# Nuclear phenotypic changes in cells originating from migrating tenocytes in tendon explants

**DOI:** 10.1101/2025.05.10.653257

**Authors:** Eli Heber Martins dos Anjos, Maria Luiza Silveira Mello, Benedicto de Campos Vidal

## Abstract

Tenocytes migrating from cultured rat tendon explants aggregate and gradually align in parallel rows with the tendon’s long axis as the explant period extends up to 12 days and collagen bundles become structurally remodeled. Tendon architecture and mechanical load are known to affect fibroblast phenotypes; thus, we studied cell cultures derived from tenocytes at early and later migration stages using image analysis and immunofluorescence for selected epigenetic markers to identify differences in nuclear characteristics across migration periods. Feulgen-DNA amounts and global DNA 5-methylcytosine and acetylated H3K9 distribution were not found to differ in the cells originating from tenocytes that migrated at the 8- and 12-day periods of tendon culture. However, light absorbance, fractal dimension, and margination features related to chromatin condensation and distribution in the Feulgen-stained cell nuclei differed during these periods. Accumulation of the H3K4me2 marker at the nuclear periphery in cells obtained after a longer explant culture period, when the margination feature reaches larger values, may represent the acquisition of an enhanced transcriptional memory that could affect future gene expression levels. Further assays are required to determine whether the present image analysis results correlate with the differential potential of these cells for extracellular matrix synthesis and practical bioengineering purposes.

## 1. Introduction

Rat tendon explants cultivated on plastic substrates exhibit structural remodeling with a reduction in macromolecular orientation and an aggregation and alignment of collagen bundles (dos Anjos et al., 2023a). This event is accompanied by tenocyte migration from the lateral zones of the tendons and their alignment with respect to the long axis of the tendons, which gradually occurs with increasing cultivation times from a 6-to 7-day period to a 12-day period, when tenocyte alignment in parallel rows reaches approximately 1000 µm distance from the explant boundary (dos Anjos et al., 2023a). The preferential cell migration from cultured rat tendons is consistent with observations by other authors of cells migrating from explants of equine digital flexor tendons and rabbit anterior cruciate ligaments (Cadby et al., 2014; Schwarz et al., 2019). The gradual alignment of tenocytes from rat tendons has been suggested to be a cellular response to biochemical signals arising from the tendon structure cultivated *in vitro*, guided by collagen signal transducer action (dos Anjos et al., 2023a).

Although cell culture models are less complex than *in vivo* models and may be limited by the loss of proportional heterogeneity from isolated cell populations during cell cultivation (Vigano et al., 2017; Wunderli et al., 2020), migrating tenocytes are considered a promising cell source for ligament reconstruction (Schwarz et al., 2019). Because the tenocyte phenotype is affected by tendon architecture and mechanical loading (Clegg et al., 2007; Vermeulen et al., 2019; Wu et al., 2022), the characteristics of the nuclear phenotypes of cells migrating from tendon explants, where the organization of collagen bundles has been found to change (dos Anjos et al., 2023a), may differ and thus constitute a matter of scientific interest for image analysis studies. Chromatin remodeling, which is frequently associated with differential accessibility for transcription factors or DNA-binding proteins (Mello, 2021), has been found to be affected by histone modifications in tendinopathic tissue (Orchard et al., 2023; Orchard, 2024).

Image analysis of Feulgen-stained nuclei has been demonstrated a useful tool to reveal changes in Feulgen-DNA amounts, nuclear phenotypes, and chromatin supraorganization in several cell models, including hepatocytes, fibroblasts, and cardiomyocytes under different physiological and/or pathological states (Moraes et al., 2005, 2007; Aldrovani et al., 2006; Ghiraldini et al., 2012; Silva et al., 2018). Image analysis features have also been proved as potential markers for detection of preneoplastic changes, tumor progression and prognosis (Adam et al., 2006; Ferro et al., 2011; Mendaçolli et al., 2015; Metze et al., 2019; Dincic et al., 2021; Onuma et al., 2024).

Loss of Con-A positive nuclear glycoproteins and an increase in chromatin packing states induced in inbred strain-A/Uni mice by food deprivation for 48 h and subsequent recovery with feeding have been assessed in hepatocytes using image analysis (Moraes et al., 2005). Hepatic polyploidy and an increase in contrast between condensed and non-condensed chromatin in Feulgen-stained preparations, assessed by image analysis, and associated with higher formation of extended chromatin fibers under gravitational force, could be detected in aged mice (Moraes et al., 2007). Significant differences in image analysis features which defined polyploid DNA amounts and nuclear phenotypes were also reported for tail tendon fibroblasts from adult non-obese diabetic (NOD) compared to healthy BALB/c mice (Aldrovani et al., 2006), and for hepatocytes from hyperglycemic NOD mice compared to normoglycemic aged BALB/c mice, thus favoring the idea of differences in chromatin functions between diabetes and aging (Ghiraldini et al., 2012). In the same context, dissimilarities in terms of ploidy degrees, chromatin texture, and nuclear shapes have been detected between cardiomyocytes from adult NOD mice (irrespective of their glycemic levels) and normoglycemic aged BALB/c mice (Silva et al., 2018).

In the present study, we used image analysis to investigate whether the nuclear image features of cells cultivated *in vitro* from tenocytes during their early period of migration from rat tendon explants (8-day period) changed after these cells acquired extensive parallel alignment with respect to the long axis of the tendon (12-day period). In addition, the nuclear epigenetic marker 5-methylcytosine (5mC) and two markers indicative of transcriptional activity, acetylated histone H3 (H3ac) and dimethylated H3K4 (H3K4me2) (Rocha et al., 2019, 2023, 2024), were investigated in these cells using confocal fluorescence microscopy.

## 2. Materials and methods

### 2.1. Ethical considerations and animals

This study was conducted in compliance with Brazilian Federal Law No. 11,794 of October 8, 2008, followed the guidelines established by the Brazilian National Council for the Control of Animal Experimentation (CONCEA), approved by the Committee of Ethics in Animal Use (CEUA/IB/UNICAMP, Campinas), and registered under protocol number 5064-1/2018. The authors declare that animal experimentation followed the ARRIVE guidelines and was carried out in accordance with the U.K. Animals (Scientific Procedures) Act, 1986, and associated guidelines. Male adult Wistar rats (*Rattus norvegicus albinus*) (n = 5) were provided by the Multidisciplinary Center of Biological Investigation (CEMIB) at the University of Campinas (Campinas, Brazil). The animals were housed in an optimal environment with controlled humidity (55 ± 10%), temperature (22 ± 2°C), photoperiod (12/12-hour light/dark cycle), light intensity, and had free access to extruded chow (Nuvital®, Colombo, Brazil) and water *ad libitum*. The animal welfare was maintained throughout the study by using humane restraint and transportation methods, handling by a single observer, daily inspections for signs of discomfort or infection, and euthanasia performed using appropriate techniques.

### 2.1. Cells, and sample preparations

Tenocytes were obtained from Achilles tendon explants cultivated *in vitro* on plastic substrates and were used to study the structural remodeling of tendon collagen bundles in a previous study (dos Anjos et al. 2023a). The mean mass of the tendon used to provide the explants was 27.5 ± 10.7 mg. The tendon explants were cultivated in DMEM/HAMF12 (1:1) (DMEM, Sigma-Aldrich^®^, Saint Louis, MO, USA; HAMF12, Vitrocell Embryolife, Campinas, Brazil), with DMEM supplemented with 1% penicillin-streptomycin (Sigma-Aldrich^®^, 100 IU/mL and 100 µg/mL, respectively), at 37 °C, with 95% humidity and 5% CO_2_ saturation for 8 and 12 days. The culture medium was replaced every 72 h. The cells that migrated from the explants and adhered to the plastic surface were transferred to plastic plates to establish a primary cell culture.

Cell viability was evaluated at the first passage of cell culture using the Trypan Blue assay. For detection of nuclear phenotype characteristics, a 2.5×10^3^ cell/mL aliquot of cells at the third cellular passage was cultured for 72 h. Tenocytes were fixed in an absolute ethanol:glacial acetic acid mixture (3:1, v/v) and subjected to the Feulgen reaction, which is specific for DNA (Mello and Vidal, 2017). For general cytochemical observations, some preparations were stained with a 0.025% toluidine blue (TB) (Merck, Darmstadt, Germany) solution in 0.1 mol.L^-1^ McIlvaine phosphate buffer at pH 4.0 for 15 min, followed by rinsing in buffer solution at pH 4.0 and treatment with a 4% ammonium molybdate (Merck, Rio de Janeiro, Brazil) aqueous solution for 4 min (Mello and Vidal, 1973). Under this staining condition, the dye molecules bind to available nucleic acid phosphate groups (Vidal and Mello, 2019).

For fluorescence assays, an aliquot of 2.5×10^3^ cells/mL, cultured for 72 h, was used, and the cells were fixed in 4% paraformaldehyde (cat no. 10,284, Lot no. 900,835; Quimibras Indústrias Químicas SA, Rio de Janeiro, RJ, Brazil) solution in phosphate-buffered saline (PBS) at pH 7.4. After fixation, the preparations were washed thrice in 70% ethanol for 1 min each and were immersed in ACS-grade methanol at -20 °C for 10 min. The slides were stored at 4-8 °C in PBS at pH 7.4 until use.

### 2.2. Image analysis

Image analysis of the Feulgen-stained nuclei was performed using an Olympus BMX-51 microscope (Olympus America Center, Center Valley, PA, USA) equipped with a Q-color 5 camera (Olympus America Center), a 40x objective, and λ = 546 nm. Image-Pro Plus v.6.3 software for Windows™ (Media Cybernetics, Inc., Silver Spring, MD, USA) was used for image capture and analysis. At least 300 nuclei selected at random were evaluated as recommended (Wied et al., 1989). The software provided quantitative information on geometric features as follows: 1) total nuclear area (µm^2^) (A_T_); 2) Feret ratio (FR), which is an indication of the elongation of the nuclei, with a value of 1.0 corresponding to a circle. The Feret ratio establishes a relation between Feret perpendicular diameters. A Feret diameter measures the distance between two parallel tangents of an object; and 3) fractal dimension (FD), an extension of conventional Euclidean geometry that informs about the degree to which the image fills the available space (Mandelbrot, 1967) or, in this case, the complexity of the nuclear contour. For the analysis of FD, the software implemented a variation of Richardson’s method. Studies of FD have been considered relevant as a potential marker reflecting the topological redistribution of chromatin and modifications of the nuclear architecture (Metze et al., 2019). The densitometric features evaluated were: 1) average gray values converted into absorbances or optical density (OD) per nucleus; and 2) integrated OD (IOD = OD x A_T_), which in the present study represents Feulgen-DNA values in arbitrary units. The textural feature chromatin margination (Pouliakis et al., 2014), which corresponds to the pattern of distribution of OD values between the center and the nuclear periphery, was also evaluated. The algorithm for the margination parameter developed by Ian Young (Delft University of Technology) indicates that a 0.33 value corresponds to a homogeneous distribution of OD values in the cell nuclei (Young et al., 1986).

### 2.3. Immunofluorescence assays

For 5mC detection, a modification of the protocol proposed by Veronezi et al. (2017) was used for HeLa cells. Initially, the preparations were immersed twice in PBS (pH 7.4). After washing, the cells were treated with 2 mol·L^-1^ HCl (cat. no. A1028.01.BJ, Lot no. 219,482; LabSynth, Diadema, SP, Brazil) for 1 h at 25° C, then washed three times in a borate buffer at pH 8.5 (100 mM boric acid (cat. no. 15583-016, Lot no. 1,019,235; Life Technologies, Grand Island, NY, USA), 75 mM NaCl (cat. no. 01021, Lot no. 20,132; Neon Commercial Ltd., Suzano, SP, Brazil), and 25 mM sodium tetraborate (cat. no. B1.020, Lot no. 32,142; LabSynth)) and subsequently blocked with 5% bovine serum albumin (BSA, cat. no. A7906, Lot no. SLCP8701; Sigma-Aldrich^®^) in phosphate buffer (pH 7.4) for 30 min. Then, the cells were incubated overnight with rabbit polyclonal anti-5mC primary antibody (1:150 dilution in 1% BSA, cat. no. IM-0812, Lot no. 21,009; RheaBiotech, Campinas, SP, Brazil) at 4 °C in the dark, followed by treatment with a goat anti-rabbit antibody conjugated to Alexa-Fluor 488 (1:500 dilution in 1% BSA; cat. no. A-11,008, Lot no. 2,284,594; Life Technologies^®^, Carlsbad, CA, USA) for 2 h in the dark, and counterstained with 4’, 6-diamidino-2-phenylindole (DAPI, cat. no. D-9542, Lot no. 079K4177; Sigma-Aldrich^®^) for 1 h in the dark at 25 °C. The preparations were then washed three times in PBS (5 min each) and mounted in VECTASHIELD^®^ (cat. no. H-1000, Lot no. V1001; Vector Laboratories, Inc., Burlingame, CA, USA).

For detection of histone markers H3K4me2 and H3K9ac, the cells were fixed in 4% PFA, washed twice in phosphate buffer at pH 7.4, permeabilized with 0.2% Triton X-100 (cat. no. T-8787; CAS no. 9002-93-1; Merck, Darmstadt, Germany) for 10 min at 25 °C, rinsed thrice in PBS (pH 7.4), and blocked with 5% BSA for 30 min at 25 °C. The cells were then incubated overnight with rabbit monoclonal anti-H3K4me2 (1:500 dilution in 1% BSA; cat. no. #9725 C64G9, Lot no. 9), and rabbit monoclonal anti-H3K9ac (1:30 dilution in 1% BSA; cat. no. #9649 C5B11, Lot no. 13) (both from Cell Signaling Technology, Inc., Danvers, MA, USA) primary antibodies at 4 °C in the dark, followed by thorough washing with PBS. To detect H3K4me2 and H3K9ac signals, the cells were treated with a goat anti-rabbit antibody conjugated to Alexa-Fluor 488 (1:500 dilution in 1% BSA; cat. no. A-11,008, Lot no. 2,284,594; Life Technologies, Carlsbad, CA, USA) for 2 h at 25 °C in the dark, and then counterstained with DAPI (cat. no. D-9542, Lot No. 079K4177; Sigma-Aldrich) for 1 h at room temperature in the dark. The preparations were then rinsed in PBS and mounted in VECTASHIELD^®^ (Vector Laboratories, Inc.). Signals for 5mC, H3K4me2, and H3K9ac were captured using a Leica TCS SP5 II confocal microscope (Leica Microsystems GmbH) at the Central Laboratory of High-Performance Technologies in Life Sciences (LaCTAD/University of Campinas).

Three independent experiments were performed to detect fluorescence intensity. The total number of nuclei analyzed was 69 and 23 for 5mC (8 and 12 days, respectively); 33 and 26 for H3K9ac (8 and 12 days, respectively); and 26 and 23 for H3K4me2 (8 and 12 days, respectively). Negative controls consisted of cells incubated only with secondary antibodies. HeLa cells (ATCC: CCL-2), acquired from the Emerging Virus Studies Laboratory (LEVE) at UNICAMP and validated at the Technical Directorate for Teaching and Research Support (DTAPEP) of the Faculty of Medicine Foundation (University of São Paulo, USP), were used as the positive control for fluorescence assays because of their previously reported response to anti-5mC and anti-histone markers H3K4me2 and H3ac antibodies (Rocha et al. 2019, 2023, 2024). An aliquot of 5.0×10□ cells/mL at passages 15-18 was cultured for 24 h, fixed in 4% PFA, and stored until use.

### 2.4. Statistics

Calculations and statistical analyses for the image analysis features were performed using Minitab 14^®^ software (State College, PA, USA) and GraphPad Prism 8.0.1 (Boston, MA, USA). The statistical analysis of fluorescence intensity was performed using GraphPad Prism 8.0.1 software. Shapiro-Wilk and Mann–Whitney tests were used to assess the statistical significance of the datasets. In all cases, a p-value of < 0.05 was considered the critical level for rejecting the null hypothesis.

## 3. Results

### 3.1. Primary cell cultures

The cell viability, determined by the ratio between the number of viable cells and the total cells that originated from tenocytes migrating from tendon explants cultured for 8 and 12 days was 100% and 94.4%, respectively. The morphological aspects of the cells and cell nuclei developed in *in vitro* cultures of tenocytes that migrated from the tendon explants are shown in Figure 1A-D. Metachromatic staining of nucleic acids was evident in tenocytes stained with TB at pH 4.0 (Fig. 1B). A comparison of the Feulgen-stained nuclei of cells originating from tenocytes that migrated from tendon explants cultured for 8 and 12 days even visually reveals phenotypic changes (Fig. 1C and D).

**Fig. 1.**
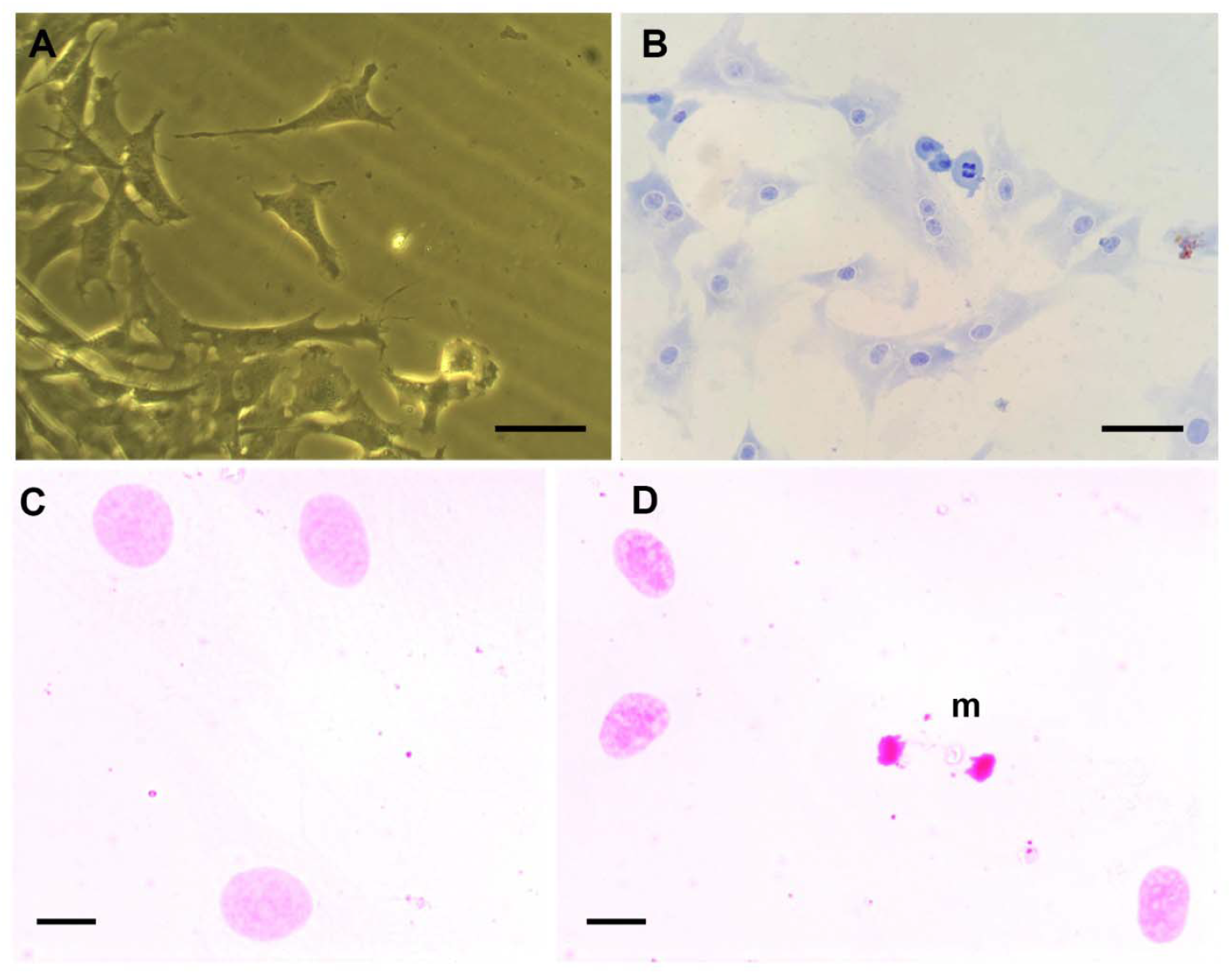
Microscopic aspects of cells derived from migrating tenocytes of tendon explants. Unstained (A) and TB-stained (B) cells are shown after the establishment of *in vitro* culture from tenocytes originally migrating from tendon explants cultivated for 12 days. The metachromatic phenomenon observed after TB staining (B) resulted from the electrostatic binding of the dye molecules to anionic groups of RNA and DNA substrates (Vidal and Mello, 2019). Images of Feulgen-stained nuclei of cells derived from tenocytes that migrated from tendon explants cultured for 8 (C) and 12 (D) days reveal a decrease in nuclear area and increased chromatin condensation with advancing tendon culture time. m, mitosis (Bars = 50 µm (C and D) and 10 µm (E and F)). TB, Toluidine blue.

### 3.2. Image analysis

The Feulgen-DNA values calculated from nuclear areas (A_T_) x OD values, or integrated OD (IOD) values (Figs. 2A-C; Tables 1, 2) were not found to change when comparing cells originated from migrating tenocytes of tendons cultured for 12 days relative to those cultured for 8 days. This result occurred because with increasing periods of tendon explant cultures the cellular nuclear areas decreased (Table 1) while OD values increased (Table 2) (Figs. 2A, B). Under these same tendon culture conditions, the nuclear Feret ratio values were not affected; they were greater than 1.0, indicating maintenance of a nuclear ellipsoidal shape (Fig. 2D; Table 1).

**Table 1.** Nuclear geometric parameters of Feulgen-stained tenocytes.

| Culture times<br>(days) | N | $A_T$ | | | FR | | | FD | | |
| --- | --- | --- | --- | --- | --- | --- | --- | --- | --- | --- |
| | | $\bar{X}$ | SD | Md | $\bar{X}$ | SD | Md | $\bar{X}$ | SD | Md |
| 8 | 305 | 644.1 | 297.2 | 632.4 | 1.402 | 0.307 | 1.344 | 1.062 | 0.051 | 1.045 |
| 12 | 319 | 511.8 | 261.4 | 472.9* | 1.398 | 0.308 | 1.356 | 1.045 | 0.030 | 1.035* |

$A_T$ , total nuclear area ( $\mu\text{m}^2$ ); FD, fractal dimension; FR, Feret ratio; Md, median; N, number of measurements; SD, standard deviation; $\bar{X}$ = arithmetic mean; \*, highly significant differences at $p < 0.01$ level (Mann-Whitney test preceded by Shapiro-Wilk test). Individual values composed a dataset that was deposited in a public repository at the University of Campinas (dos Anjos et al., 2023b).

**Table 2.** Nuclear densitometric and textural parameters of Feulgen-stained tenocytes.

| Culture times (days) | N | OD |  |  | IOD |  |  | Margination |  |  |
| --- | --- | --- | --- | --- | --- | --- | --- | --- | --- | --- |
| | | $\bar{X}$ | SD | Md | $\bar{X}$ | SD | Md | $\bar{X}$ | SD | Md |
| 8 | 305 | 0.2145 | 0.0561 | 0.2096 | 130.3 | 58.37 | 127.0 | 0.3210 | 0.0157 | 0.3248 |
| 12 | 319 | 0.2540 | 0.0423 | 0.2416* | 124.5 | 56.28 | 110.8 | 0.3164 | 0.0150 | 0.3193* |

IOD, integrated optical density (Feulgen-DNA values in arbitrary units); Md, median; N, number of measurements; OD, optical density; SD, standard deviation; $\bar{X}$ = arithmetic mean; \*, highly significant difference at the $p < 0.01$ level. Individual values composed a dataset that was deposited in a public repository at the University of Campinas (dos Anjos et al., 2023b).

**Fig. 2.**
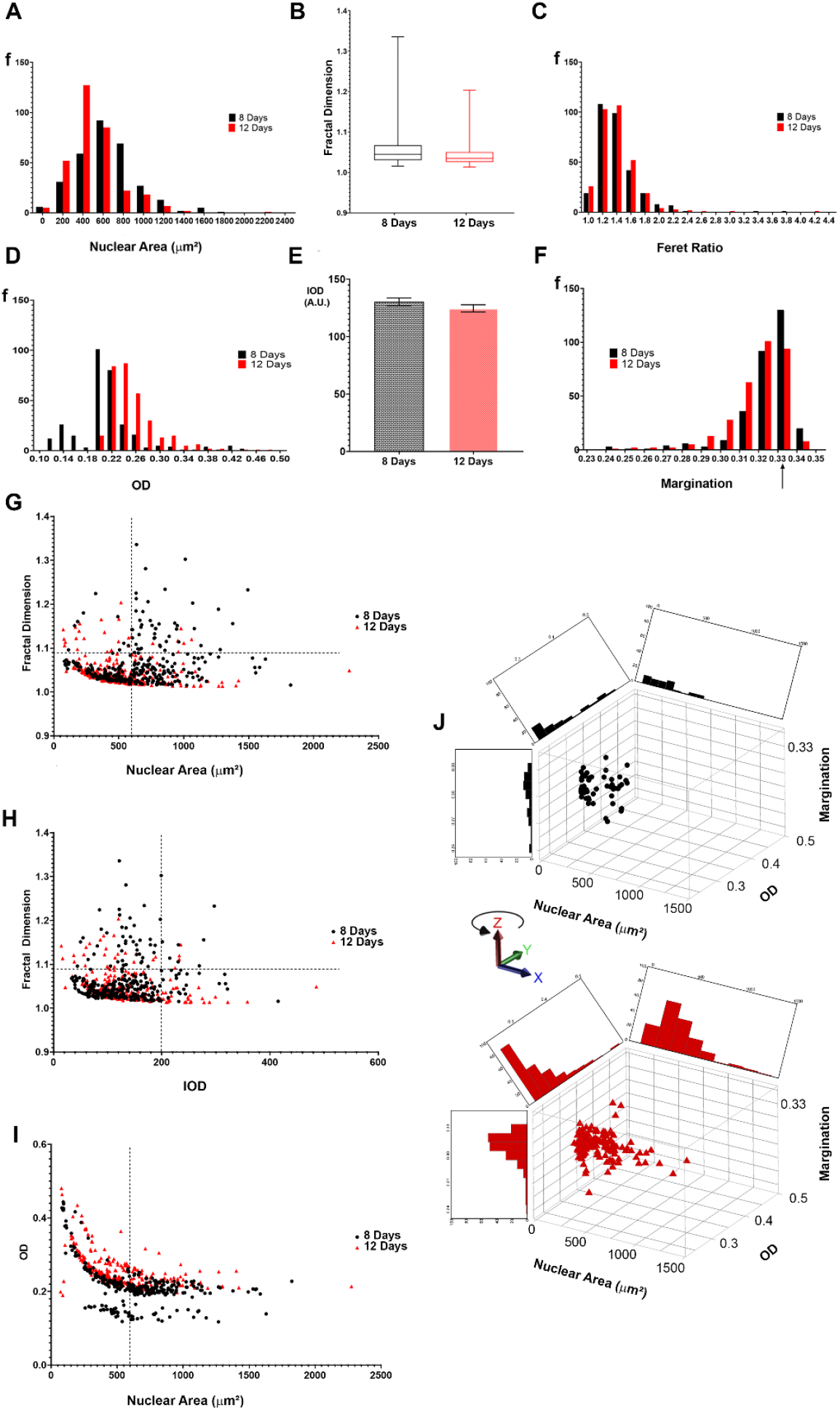
Image analysis features of Feulgen-stained cells derived from migrating tenocytes of tendon explants under different tendon culture times. Frequency histograms (A-D, F), a box- and-whisker plot (E), and relation between features (G-J). The value 0.33 on the X-axis for the margination feature (F) refers to nuclei containing a homogeneous distribution of chromatin OD values (Young et al., 1986). The distribution of points corresponding to individual cell nuclei whose feature values change as a function of the original tendon culture times is evident (G-J). A.U., arbitrary units; f, frequency; IOD, Feulgen-DNA values; OD, absorbances; X, Y, Z, coordinate axes

A significant decrease in FD values, reflecting changes in the nuclear architecture, and a shift of the distribution of the nuclear margination data from value 0.33 occurred in the cells originated from the tenocytes which migrated from the tendon explants cultured longer (Figs. 2E, F; Tables 1, 2).

Most nuclei with areas >/ 600 µm^2^ and FD values >/ 1.09 in cells originated from tenocytes which migrated from 8-day tendon explants, subsequently experienced a decrease in these values (Fig. 2G), indicating a reduced complexity in chromatin occupancy under the longer period of tendon culture. However, a correlation between FD and IOD values did not reveal differences when comparing cells obtained from the two periods of tendon culture. Nuclei with IOD values < 200.0 and FD < 1.09 represented 75.1% and 82.1% of cells cultured from tenocytes migrating from tendon explants cultured for 8 and 12 days, respectively (Fig. 2H).

Regardless of nuclear size, higher OD values were detected in cells originating from tenocytes that migrated when tendon explant cultivation was extended to 12 days (Fig. 2I). Margination values < 0.33, which were linked with increased OD values and decreased nuclear sizes, were observed in cells originating from tenocytes migrating from tendon explants cultured for 12 days (Fig. 2J).

### 3.3. Immunofluorescence

No change in the fluorescence intensity of 5mC and H3K9ac signals was detected when comparing cells originating from migrating tenocytes of tendon explants cultured for 8 and 12 days (Fig. 3A, B, E and F). However, for the fluorescence signals for H3K4me2, a significant increase in fluorescence intensity was revealed in cells originating from tendon explants cultured for 12 days (Fig. 3C, D and G). In these cells, most nuclei exhibited intense fluorescence signals near the nuclear periphery (Fig. 3D).

**Fig. 3.**
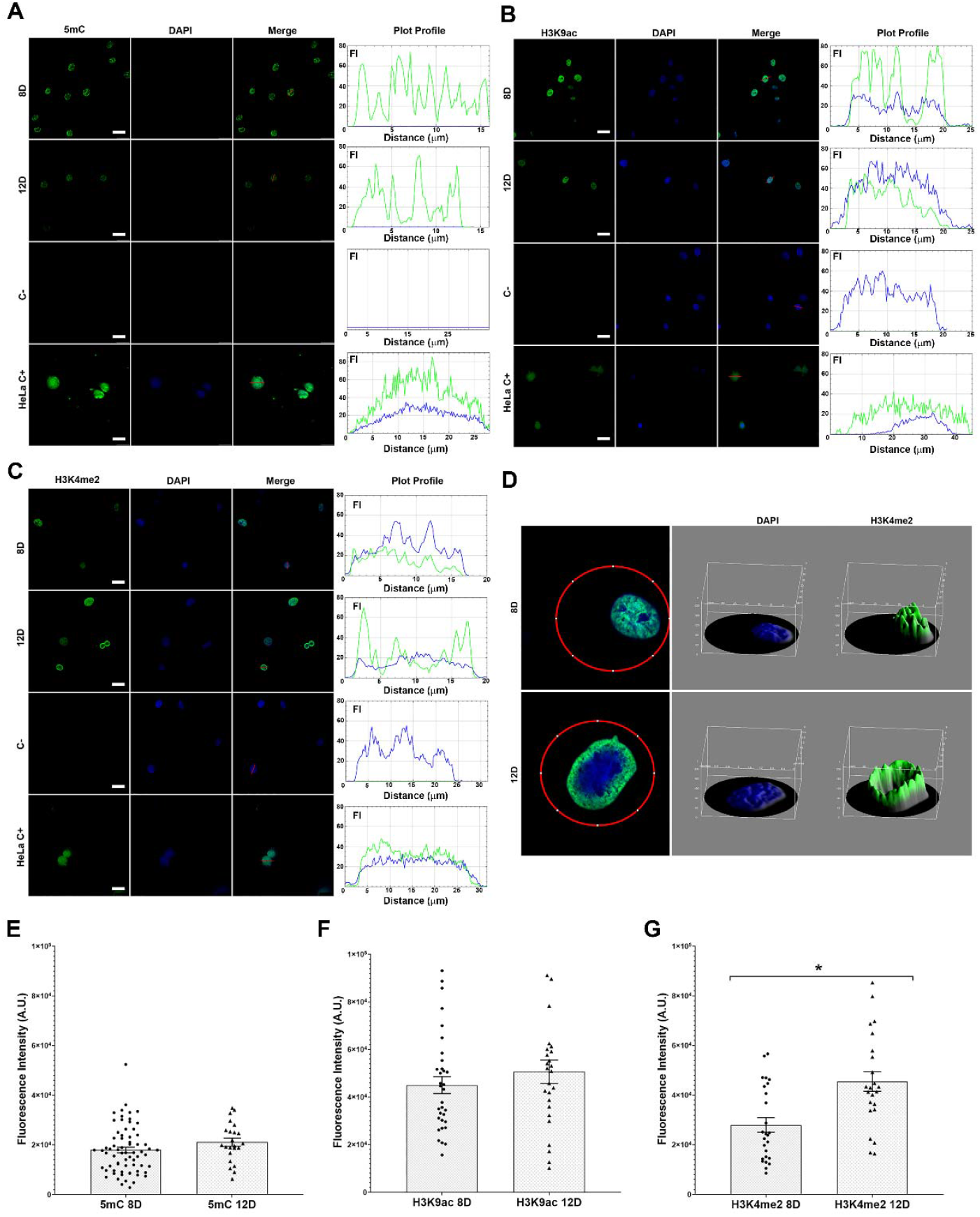
Immunofluorescence intensity of 5mC, H3K9ac, and H3K4me2 signals in cells derived from migrating tenocytes analyzed using confocal microscopy. Images A-C are representative of three independent experiments. The total number of nuclei analyzed was 69 and 23 for 5mC (8- and 12-day tendon culture times, respectively), 33 and 26 for H3K9ac (8- and 12-day tendon culture times, respectively), and 26 and 23 for H3K4me2 (8- and 12-day tendon culture times, respectively). HeLa cells were used as positive controls. Bars = 25 µm (A: 5D, 12D, HeLa C+; B: 8D, 12D, C-; C: 8D, 12D, C-, HeLa C+) and 50 µm A: C-; B: HeLa C+). A selectively more intense fluorescence response for H3K4me2 at the nuclear periphery was evident in most nuclei of the cells derived from tendons cultured longer (D). The Mann-Whitney test was used for fluorescence intensity (FI) analysis (E-G). A significant difference in H3K4me2 signals between cells resulting from 8- and 12-day periods of cultured tendon explants at P<0.05 is indicated (G) (*). The vertical lines over the graphic columns represent the standard error of the mean. C-, negative control; FI, fluorescence intensity in arbitrary units (A.U.).

## 4. Discussion

Present image analysis data indicate that the Feulgen-DNA amounts in the cell cultures originating from the tenocytes that migrated after early and longer periods of the tendon explant culture remained constant. However, features related to nuclear sizes, and to chromatin condensation and distribution patterns were found to differ. The decrease in nuclear size and FD values in cells derived from tenocytes that migrated at the 12-day of the tendon explant culture indicates that the chromatin acquired a less complex topological occupancy and suffered reduction in its accessibility at this time (Metze et al., 2019). Changes in FD reflect topological chromatin redistribution and modifications in nuclear architecture (Metze et al., 2019). The increased values of the margination feature, associated with high absorbance values and small nuclear sizes in a portion of the cell sample derived from tenocytes obtained after the 12-day long tendon culture period, aligned with the FD data.

An elongated nuclear shape persisted in cultured cells originating from both periods of tenocytes’ migration from the tendon explants. However, this shape was not as fusiform and slender as that observed in the tenocytes aligned in straight rows along the longitudinal axis of tendons cultured for 12 days (dos Anjos et al., 2023a) or in fibroblasts of adult Achilles tendons. Such a loss of the original spindle-shaped characteristics of tenocytes has been previously reported for cells isolated at different passages from rat tendon tissues (Vermeulen et al., 2019).

Although the abundance and distribution of global DNA 5mC-positive signals in the cells originating from tenocytes that migrated after earlier and longer periods of tendon explant cultures were not found to differ, this does not exclude the possibility of differences in DNA CpG methylation affecting selected genes as the tendon explant periods advances. Differential methylation at the promoters of *Leprel2, Foxf1, Mmp25, Igfbp6*, and *Peg12* genes, for instance, has been associated with fiber disorganization in murine Achilles tendons (Trella et al., 2017). DNA CpG methylation is implicated in the activity of certain fibrosis-associated genes and transcription factors in fibroblasts (Ulukan et al., 2019) although not in growth differentiation factors GDF5, GDF6, and GDF7 in tenocytes during rotator cuff tendinopathy (Orchard, 2024).

While no change occurred in the overall fluorescence signals for the H3K9ac, a marker correlated with transcriptionally active chromatin (Sterner and Berger, 2000), in the cells derived from tenocytes obtained after 8- or 12-day periods of tendon explant culture, H3K4me2 signals appeared more closely positioned at the nuclear periphery in cells obtained after the older period of the explant culture. These are the cells which exhibit larger margination values in Feulgen-stained preparations. The H3K4me2 marker in mammalian cells potentially reflects distinct regulatory mechanisms that finely tune gene expression (Pekowska et al., 2010) and is associated with a small subset of genes rapidly activated upon stimulation (Russ et al., 2014). In addition, the accumulation of H3K4me2 markers at the nuclear periphery of different cell types has been associated with enhanced transcriptional memory that can last for several days, thus influencing future expression levels (Light and Brickner, 2013; Fiserová et al., 2017). In HeLa cells, deep fluorescence intensity of H3K4me2 signals at the nuclear periphery is elicited in the presence of sodium valproate, a histone deacetylase inhibitor (Rocha et al., 2023). In the context of cells generated from tenocytes that migrated from tendon explants, it is possible that by advancing the period of the explant culture up to 12 days, enhanced transcriptional memory from specific genes spatially organized at the nuclear periphery may have been acquired. Studies on other histone marks, in conjunction with transcriptional activity data, are required for a better interpretation of these data.

Because the tendon explant culture conditions originally developed to induce the migratory activity of tenocytes did not exceed 12 days, providing cells with a homogenous distribution, increased cell numbers, and morphology like that of tendon fibroblasts (dos Anjos et al., 2023a), these conditions were considered adequate for the present study. High and consistent cell viability suggest that the culture conditions were appropriate and did not impact cell physiology.

Whether the cells that presented differences in image analysis characteristics, provided by either the original 8-day or 12-day explant culture, have a differential capacity for synthesis of the extracellular matrix (ECM), requires further investigation that will include immunolabeling, detection of cytoplasmic markers, and gene expression assays (Schwarz et al., 2019). Evaluation of the expression of the tendon markers scleraxis, fibroblast marker tenascin C, type I and III collagen, decorin, and glycoprotein lubricin is required to determine the capacity of cells to produce ECM (Yao et al., 2006; Schwarz et al., 2019; Wunderli et al., 2020). In addition, maintaining the differentiation potential of these cells to self-assemble into spheroids and emigrate into polyglycolic acid scaffold culture formations is considered essential for comparison with cells *in situ*, particularly if intended for tissue engineering purposes (Schwarz et al., 2019).

## 5. Conclusion

Image analysis of the cells derived from tenocytes that migrated earlier from rat tendon explants demonstrates no change in Feulgen-DNA amounts but changes in chromatin supraorganization and distribution patterns in cell nuclei at a longer explant culture period. In cells obtained under the longer culture period analyzed, increased values of the nuclear margination feature evaluated in Feulgen-stained preparations, and the accumulation of the H3K4me2 marker at the nuclear periphery, as assessed by immunofluorescence, may be related to the acquisition of a transcriptional memory whose effects extend to future gene expression levels. Before intending whether the image analysis data in cells derived from different periods of tenocyte migration from explants support the selection of cells for tissue engineering, it is important to test the expression of differentiation markers in the chromatin of these cells and to monitor their capacity for sufficient ECM synthesis, self-assembly, and scaffold culture formations.

## Acknowledgments

The authors are indebted to José Luiz Módena and Cristina P. Vicente for the facilities at their laboratories (UNICAMP, Campinas, Brazil), to Camila Borges M. de Oliveira for her expert assistance with cell cultures, and to Mateus Modin (USP, Piracicaba, Brazil) for his assistance with 5mC immunofluorescence assay.

## CRediT authorship contribution statement

B.C.V. and M.L.S.M. conceived and designed the experiments, contributed the reagents/materials/analysis tools, and supervised the study. E.H.M.d.A. and B.C.V. performed the experiments. E.H.M.d.A., B.C.V. and M.L.S.M. analyzed the data. M.L.S.M. and E.H.M.d.A. wrote the original draft of the manuscript. All authors have read and agreed to the published version of the manuscript.

## Informed consent statement

Not applicable.

## Data availability statement

Raw image analysis feature dataset is deposited in the repository of the University of Campinas at https://doi.org/10.25824/redu/7OQWES

